# Metadichol induces CD14 Glycoprotein Expression in Human Embryonic Stem Cells and Fibroblasts

**DOI:** 10.1101/2024.08.26.609762

**Authors:** P. R. Raghavan

## Abstract

CD14, or cluster of differentiation 14, is a glycoprotein essential to the immune system and is found primarily on monocytes, macrophages, and other immune cells. Despite its importance, there are no examples in the literature of small compounds that can induce multifold expression of CD14 in human embryonic stem cells (hESCs) or fibroblasts. This study addresses this gap by exploring the potential of metadichol, a nanoemulsion of long-chain fatty alcohols, to induce CD14 expression in hESCs. Using quantitative real-time PCR (qRT□PCR) and Western blotting techniques, we showed that metadichol significantly upregulated CD14 expression by seventeen -fold in hESCs but downregulated it in fibroblasts. This novel finding indicates that metadichol can modulate CD14 expression in a cell type-specific manner, highlighting its potential for enhancing stem cell-based therapeutics and advancing our understanding of stem cell biology. The implications of these findings are substantial, suggesting new directions for research into the immune modulatory functions of hESCs and their potential applications in regenerative medicine. Our work highlights the potential of metadichol as a powerful tool for modulating CD14 expression in stem as well as somatic cells marking a significant step forward in the field of stem cell research and therapeutic development.

## Introduction

The CD14 glycoprotein, a coreceptor for toll-like receptors (TLRs), particularly TLR4, is essential in the innate immune response because it recognizes lipopolysaccharides (LPSs) from bacterial cell walls (1).. This interaction significantly enhances the immune response to bacterial infections, making CD14 a key component in pathogen recognition and immune activation. However, the expression and functional implications of CD14 in human embryonic stem cells (hESCs) remain underexplored, revealing a unique opportunity for novel therapeutic interventions.

Inducing the expression of CD14 in human embryonic stem cells (hESCs) and fibroblasts is challenging due to several factors related to the nature and regulation of CD14 expression. In terms of tissue distribution, CD14 is predominantly expressed in monocytes and macrophages, but it is also present in dendritic cells and, to a lesser extent, in neutrophils (2). The expression of CD14 is tightly regulated by specific transcription factors and signaling molecules that are present in myeloid cells but may be absent or inactive in hESCs and fibroblasts. For example, stimuli such as LPS, dimethyl sulfoxide (DMSO), and 1,25-dihydroxyvitamin D3 can induce CD14 expression in promonocytic cell lines such as U937 and HL-60, but these factors may not have the same effect on hESCs or fibroblasts because of differences in receptor expression and intracellular signaling pathways (3).

In some non myeloid cells, CD14 can be present in a soluble form (sCD14), which can participate in signaling pathways indirectly by interacting with other receptors, such as Toll-like receptor 4 (TLR4)(4). However, the mechanisms and conditions under which sCD14 influences CD14 expression or function in hESCs and fibroblasts are not well understood.

Overall, the difficulty in inducing CD14 expression in hESCs and fibroblasts is largely due to the specialized role of CD14 in immune cells and the lack of necessary transcriptional and signaling components in these nonmyeloid cell types. Metadichol, a nanoemulsion of long-chain alcohols, has shown promising effects on stem cells. Additionally, the antioxidant and anti-inflammatory properties of metadichol further support its beneficial effects on stem cells and their microenvironment. their regenerative potential.

The search results do not provide specific evidence that small molecules that increase CD14 expression are multifold in stem cells. However, they offer some insights into the expression and regulation of CD14 in different contexts:

1. CD14 is not typically a marker associated with most stem cells, such as mesenchymal stem cells (MSCs), which are generally characterized by low or absent CD14 expression (5). However, CD14 expression has been observed in certain progenitor or stem-like cells, such as porcine spermatogonial stem cells (SSCs), where it is associated with stemness genes such as OCT4 and NANOG (6).
2. Small molecules are used to influence stem cell characteristics and differentiation potential. For example, certain small molecules can affect gene expression patterns and enhance differentiation efficiency in amniotic fluid stem cells (AFSCs), although these studies did not specifically focus on CD14 expression (7).
3. In myeloid cells, small molecules that can either increase or decrease CD14 expression during differetiation processes have been identified. For example, certain inhibitors can prevent CD14 upregulation in monocytes during macrophage differentiation (8). However, this research focused more on immune cells than stem cells.

In summary, while small molecules can modulate gene expression and differentiation in stem cells, there is no direct evidence in the literature indicating that small molecules specifically increase CD14 expression in multiple types of stem cells. In this study, via Q-RT□PCR and western blotting, we demonstrated that metadichol (9) treatment significantly increased CD14 expression in hESCs. These findings suggest that metadichol modulates signaling pathways that regulate CD14 expression, potentially enhancing the immune modulatory functions and differentiation potential of hESCs.

### Experimental

All work was outsourced commercially to the service provider Skanda Life Sciences Pvt Ltd., Bangalore, India. The primers used were obtained from Saha BioSciences, Hyderabad, India, and the antibodies used were from Elabscience®, Houston, Texas, USA. hESC BG01V was obtained from ATCC®.

**Table 1:**
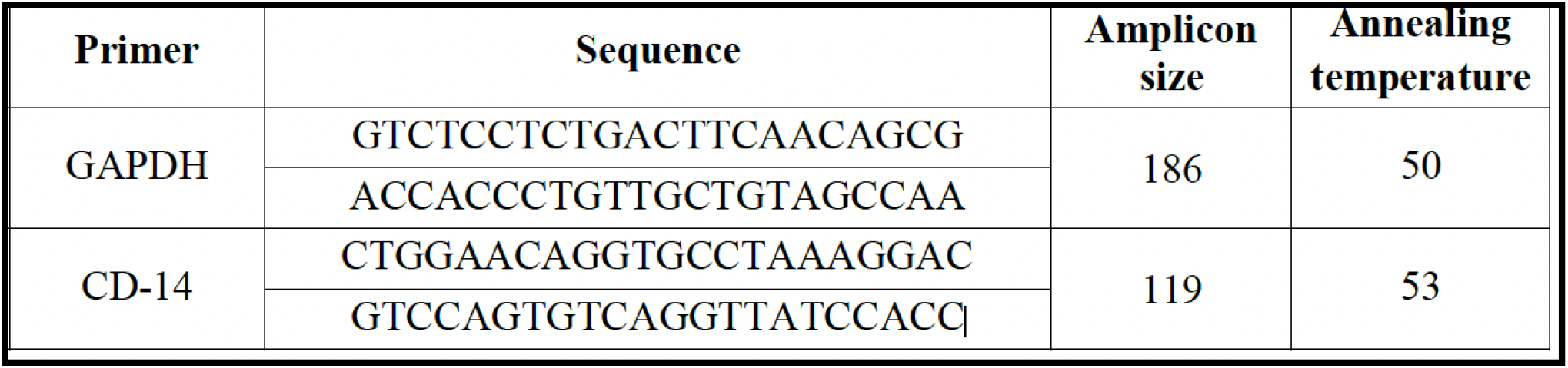
Primer details.

### Cell Culture

Human embryonic stem cells (hESCs) were maintained in suitable media with or without specific supplements and 1% antibiotics in a humidified atmosphere of 5% CO2 at 37 °C. The media was changed every other day until the cells reached confluency. Cell viability was assessed via a hemocytometer. At 70–80% confluency, a single-cell suspension at a density of 10^6 cells/mL of media was prepared and seeded into 6-well plates at a density of 1 million cells per well. The cells were incubated for 24 hours at 37 °C with 5% CO2. After 24 hours, the cell monolayer was rinsed with serum-free media and treated with predetermined concentrations of Metadichol.

### Cell Treatment

Metadichol was prepared at various concentrations (1 pg/mL, 100 pg/mL, 1 ng/mL, and 100 ng/mL) in serum-free media and added to the respective predesignated wells. The control cells received only media without the test sample. The cells were incubated for 24 hours, after which the regulation of various biomarkers was analyzed via quantitative real-time PCR (qRT□PCR) and Western blot techniques.

### Quantitative real-time PCR

Total RNA was isolated from each treatment group via TRIzol Reagent (Invitrogen, Carlsbad, CA, USA). The cells at a density of approximately 10^6 cells were collected in 1.5 mL microcentrifuge tubes and centrifuged at 5000 rpm for 5 minutes at 4 °C. The cell supernatant was discarded, and 650 µL of TRIzol was added to the pellet. The contents were mixed well and incubated on ice for 20 minutes. To the mixture, 300 µL of chloroform was added, and the mixture was mixed well by gentle inversion for 1–2 minutes, followed by incubation on ice for 10 minutes. The contents were subsequently centrifuged at 12000 rpm for 15 minutes at 4 °C. The upper aqueous layer was carefully transferred to a new sterile 1.5 mL centrifuge tube, and an equal amount of prechilled isopropanol was added. The mixture was incubated at -20 °C for 60 minutes and then centrifuged at 12000 rpm for 15 minutes at 4 °C. The supernatant was discarded, and the RNA pellet, which was washed with 1.0 mL of 100% ethanol followed by 700 µL of 70% ethanol under the same centrifuge conditions, was retained. The RNA pellet was air-dried at room temperature for approximately 15–20 minutes and then resuspended in 30 µL of DEPC-treated water. The RNA was quantified via a SpectraDrop system (Molecular Devices, San Jose, CA, USA), and cDNA synthesis was carried out via reverse transcriptase PCR.

cDNA was synthesized from 2 µg of RNA via the PrimeScript RT reagent kit (TAKARA, Shiga, Japan) with oligo dT primers according to the manufacturer’s instructions. The reaction volume was set to 20 µL, and cDNA synthesis was performed at 50 °C for 30 minutes, followed by RT inactivation at 85 °C for 5 minutes via a Veriti Thermal Cycler (Applied Biosystems, Foster City, CA, USA). The cDNA was further used for real-time PCR analysis.

The PCR mixture (final volume of 20 µL) contained 1 µL of cDNA, 10 µL of SyBr Green Master Mix (Thermo Fisher Scientific, Waltham, MA, USA), and 1 µM of the respective complementary forward and reverse primers specific for the target genes. The samples were initially denatured at 95 °C for 5 minutes, followed by 30 cycles of secondary denaturation at 95 °C for 30 seconds, annealing for 30 seconds at the optimized temperature, and extension at 72 °C for 1 minute. The optimal number of cycles was selected to ensure that the amplifications were in the exponential range and did not reach a plateau. The results were analyzed via CFX Maestro Software (Bio-Rad, Hercules, CA, USA).

### Protein isolation

Total cellular protein was isolated from 10^6 cells via RIPA buffer supplemented with PMSF protease inhibitor (Sigma□Aldrich, St. Louis, MO, USA). The cells were lysed for 30 minutes at 4 °C by gentle inversion. The lysate was centrifuged at 10000 rpm for 15 minutes, and the supernatant was transferred to a fresh tube. The protein concentration was determined via the Bradford method (Bio-Rad, Hercules, CA, USA). A total of 25 µg of protein was loaded into a gel with 1X sample loading dye containing SDS. Proteins were separated under denaturing conditions using Tris□glycine running buffer.

### Western blot

Proteins were transferred to methanol-activated PVDF membranes (Invitrogen, Carlsbad, CA, USA) via the Turbo Trans-Blot system (Bio-Rad, Hercules, CA, USA). The membranes were blocked with 5% BSA for 1 hour and incubated with the appropriate primary antibodies overnight at 4 °C. This was followed by incubation with a species-specific secondary antibody for 1 hour at room temperature. The blots were washed and incubated with an enhanced chemiluminescence (ECL) substrate (Merck, Darmstadt, Germany) for 1 min in the dark. Images were captured with appropriate exposure using the ChemiDoc XRS system (Bio-Rad, Hercules, CA, USA).

## Results

**Figure 1:**
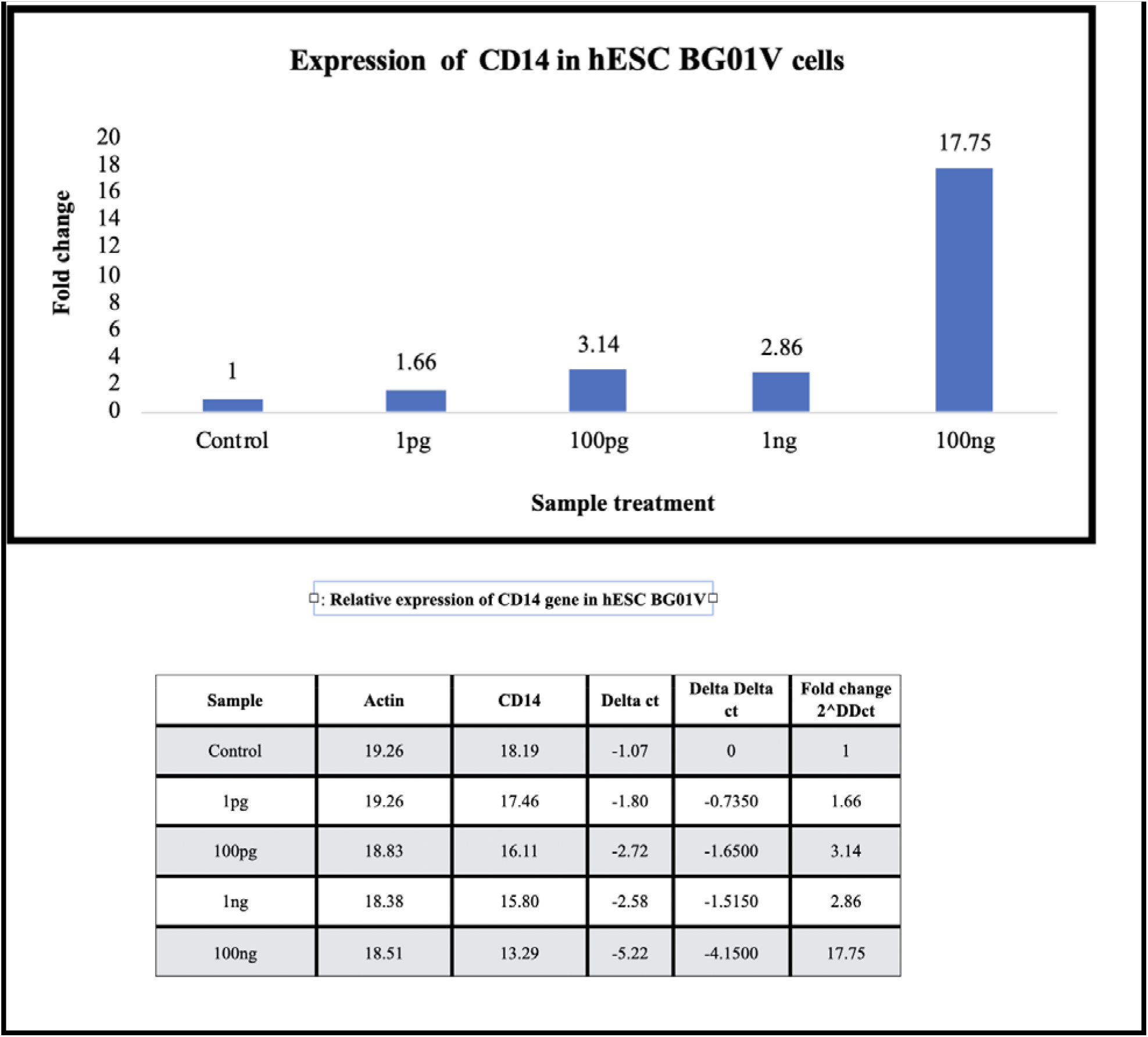
Q-RT-PCR. CD14-fold regulation in h-ESC

**Figure 2;.**
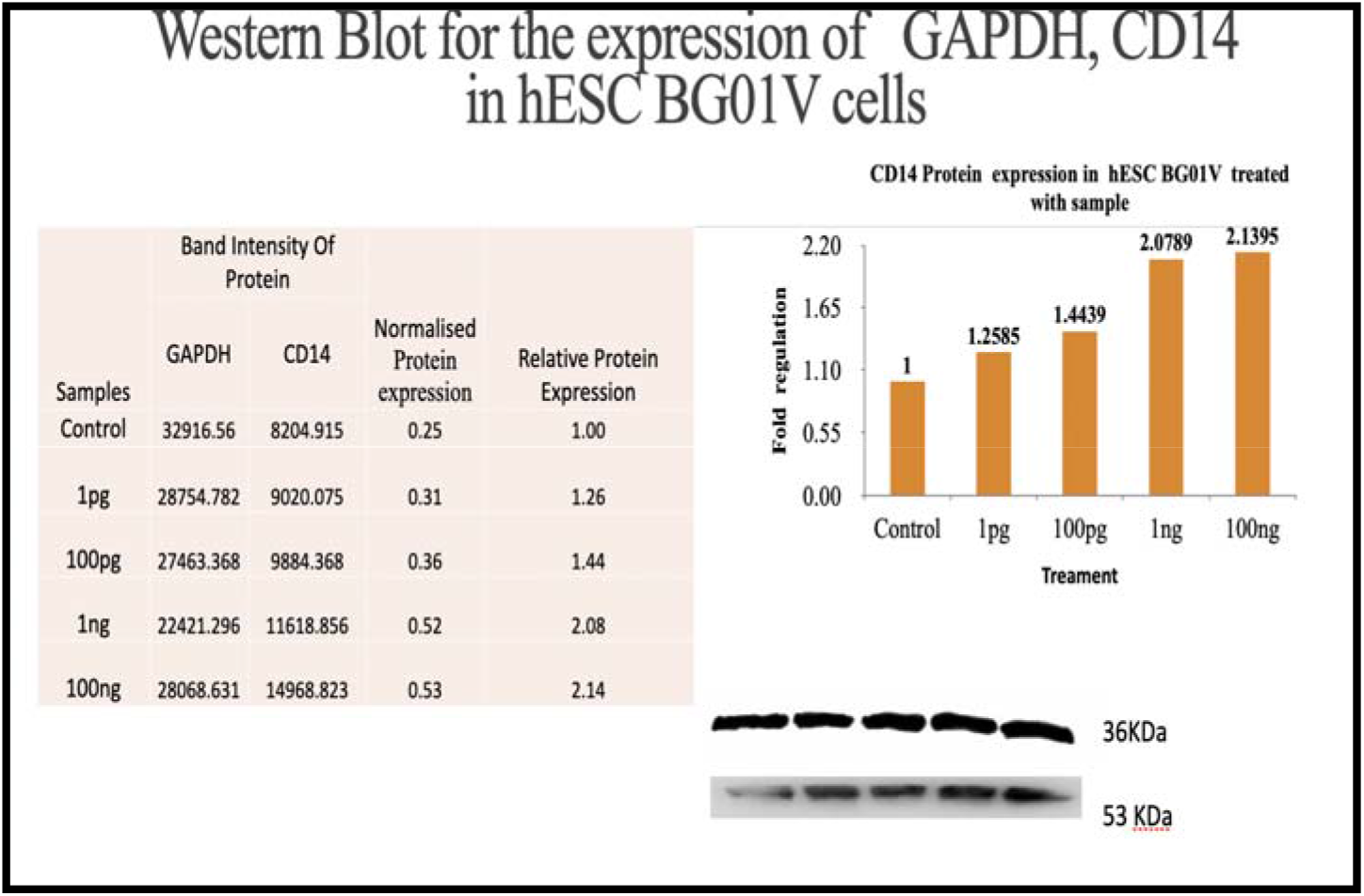
Western blot CD14 protein.

**Figure 3;.**
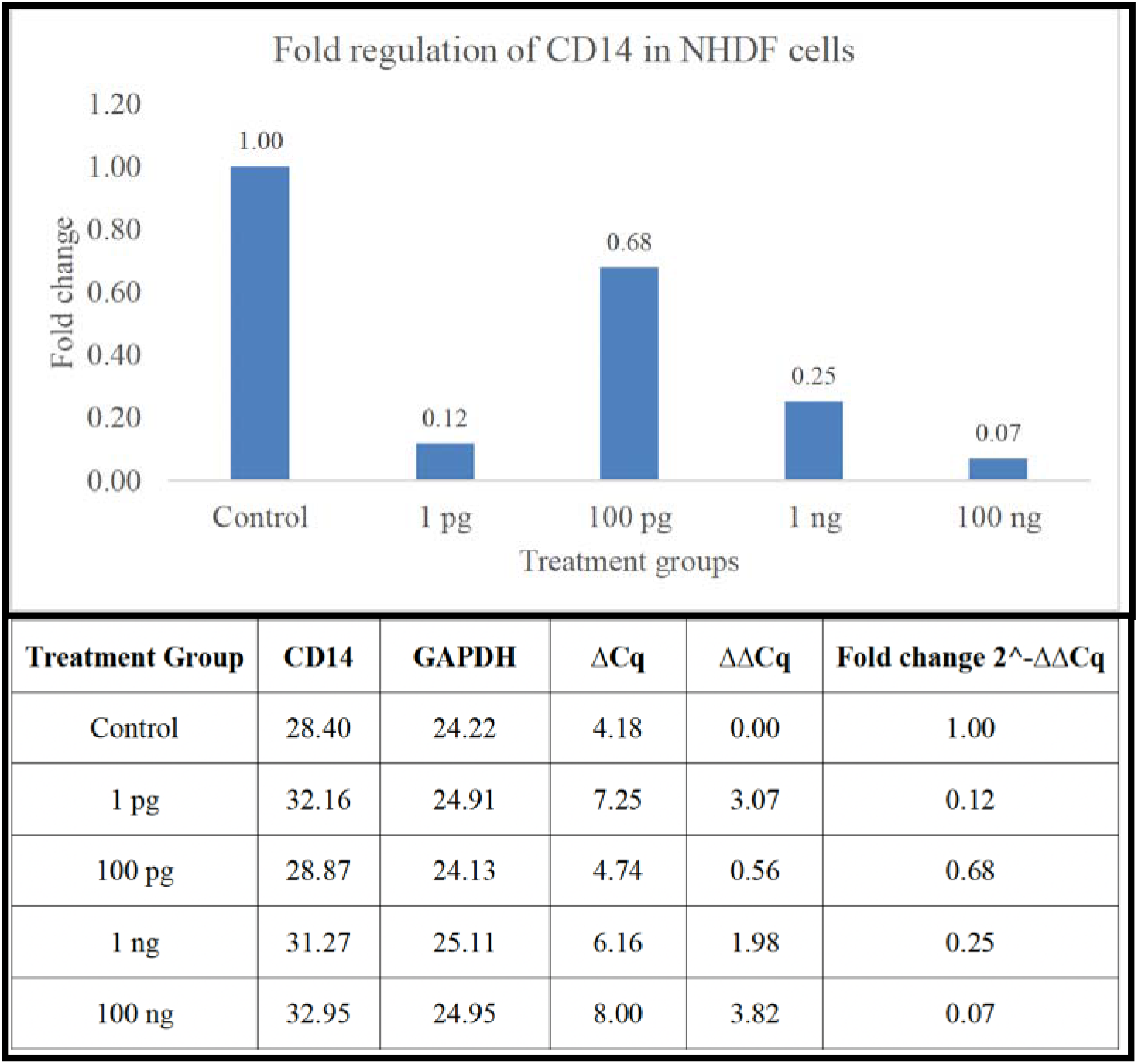
Q-RT-PCR; CD 14 fold regulation in NHDF cells.

Compared with the literature, we observed that while CD14 expression has been studied in various cell types, including MSCs and hematopoietic stem cells, there is no information on its expression in hESCs. Previous studies have shown that CD14 expression can be induced in different cell types under various conditions (10). Our study adds to this body of knowledge by demonstrating that metadichol can induce CD14 expression in hESCs, a novel finding that has not been previously reported.

## Discussion

The effects of metadichol on CD14 expression in different cell types are multifaceted and can be understood in the context of its potential applications in regenerative medicine, cancer therapy, and cellular reprogramming (11).

The ability of metadichol to increase CD14 expression seventeen-fold in h-ESCs suggests a significant impact on the immune-modulatory functions of these cells. CD14 is a coreceptor for the detection of bacterial lipopolysaccharides and plays a role in the innate immune response. The increased expression of these genes could increase the ability of stem cells to modulate immune responses, potentially improving their therapeutic efficacy in regenerative medicine and immune-related disorders (12).

The complete downregulation of CD14 in fibroblasts by Metadichol could imply a reduction in the inflammatory response typically mediated by these cells. Fibroblasts are involved in wound healing and tissue repair, and their role in inflammation is crucial. By downregulating CD14, Metadichol might reduce chronic inflammation and fibrosis, which could be beneficial in treating fibrotic diseases and improving tissue regeneration (13,14).

Nuclear hormone receptors (NHRs) play a significant role in modulating CD14 expression, primarily through their ability to regulate immune and inflammatory responses. Several NHRs are involved in this regulatory process.

PPARs (peroxisome proliferator-activated receptors), particularly PPARγ, are involved in modulating inflammatory responses and have been shown to influence the expression of CD14. PPARγ is associated with alternative (M2) macrophage polarization, which is linked to anti-inflammatory responses (15)

VDRs are known to play a role in immune regulation. The activation of VDRs can influence the expression of CD14, as vitamin D is involved in modulating the immune response and inflammation(16).

The search results indicate that the link between the vitamin D receptor (VDR) and CD14 involves at least one primary pathway, which is mediated through the interaction of the VDR with PI 3-kinase signaling. The PI 3-kinase pathway is crucial for the D3-induced expression of CD14, even though the CD14 promoter does not contain a canonical vitamin D response element (VDRE) (17).

While direct binding of the VDR to the CD14 promoter is not evident, the VDR may influence CD14 expression through other signaling molecules or transcription factors that that interact with the CD14 promoter. The expression of CD14 is tightly regulated at the transcriptional level, and Sp1 is a critical transcription factor involved in this regulation. Sp1 binds to specific regions in the CD14 promoter, and its binding is essential for the tissue-specific expression of CD14 in monocytic cells (18-20). Sp1 can also influence the expression of the VDR gene itself. The promoter region of the VDR gene contains Sp1 binding sites, which are essential for the transcriptional activation of VDR. This suggests that Sp1 not only partners with VDR to regulate other genes but also plays a role in modulating VDR expression levels. Metadichol is an inverse agonist more likely a protean agonist (21,22) of the VDR (9)

Role in immune cells. VDR is known to regulate CD14 expression in immune cells, such as macrophages, where it is a target of vitamin D signaling. In these cells, vitamin D can upregulate CD14 expression, which is important for immune responses (23).

RXRs form heterodimers with other NHRs, such as PPARs and LXRs, enhancing their ability to regulate gene expression, including that of CD14 (224. These nuclear receptors interact with various signaling pathways to regulate immune responses, and their activation by specific ligands can potentially lead to changes in CD14 expression. This modulation is crucial for maintaining immune homeostasis and can have therapeutic implications in conditions characterized by immune dysregulation.

Metadichol expresses all the nuclear receptors that play a role in Cd14 expression. The results of our previous study (25) are shown in Table 3. It is likely that CD14 is expressed though the activation of multiple NHRs and the interaction of VDR and SP1, which play a role in CD14 regulation. Further work is underway to elucidate the exact mechanism involved.

**Table 3.**

| **Fibroblasts** | **Control** | **1 pg** | **100 pg** | **1 ng** | **100 ng** |
| --- | --- | --- | --- | --- | --- |
| PPARG | 1 | 3.78 | 6.11 | 7.31 | 3.07 |
| VDR | 1 | 1.83 | 2.34 | 2.35 | 1.54 |
| RXra | 1 | 2.5 | 0.86 | 1.32 | 0.98 |
| RXRb | 1 | 4.21 | 1.65 | 1.03 | 2.7 |
| RXR g |  | 2.84 | 2.95 | 3.9 | 1.09 |
| Stem cells | **1** | **1 pg** | **100 pg** | **1 ng** | **100 ng** |
| PPARG | 1 | 1.82 | 1.7 | 1.03 | 1 |
| VDR | 1 | 2.03 | 0.92 | 3.67 | 0.54 |
| RXra | 1 | 1.4 | 1.21 | 0.99 | 0.79 |
| RXRb | 1 | 1.87 | 1.13 | 1.05 | 0.69 |
| RXR g | 1 | 2.15 | 2.2 | 1.5 | 0.76 |

## Conclusions

The findings from studies on the promotion of CD14 glycoprotein expression by Metadichol in human embryonic stem cells (hESCs) and other somatic cells have several significant implications.

### Advancements in regenerative medicine

The ability of metadichol to upregulate CD14 expression in hESCs suggests potential applications in regenerative medicine. By enhancing the immune-modulatory functions of stem cells, metadichol could improve the therapeutic efficacy of stem cell-based therapies, potentially leading to more effective treatments for various diseases and injuries.

### Potential in Cancer Therapy

The modulation of CD14 expression in cancer cells by Metadichol could have implications for cancer therapy. CD14 is associated with inflammatory and proliferative tumor microenvironments. Altering CD14 expression might increase the susceptibility of cancer cells to immune surveillance or therapeutic interventions, suggesting a novel approach for cancer treatment.

### Reduction in Inflammation and Fibrosis

In fibroblasts, metadichol downregulates CD14 expression, which can reduce chronic inflammation and fibrosis. These findings suggest potential therapeutic applications for treating fibrotic diseases and improving tissue regeneration by mitigating excessive inflammatory responses.

### Insights into Stem Cell Biology

This study provides new insights into the expression and regulation of CD14 in hESCs, a previously underexplored area. This could lead to a better understanding of stem cell biology and the development of novel therapeutic strategies that leverage the immune-modulatory capabilities of stem cells.

### Mechanistic Understanding of CD14 Regulation

These findings suggest that metadichol may influence CD14 expression through interactions with nuclear hormone receptors (NHRs), such as PPARs and VDRs, which are involved in immune and inflammatory responses. Understanding these mechanisms could provide new targets for drug development and therapeutic interventions.

## Declarations

Access to data and materials, The manuscript contain all raw data.

## Conflicting goals

Author is founder and main shareholder of Nanorx, Inc., NY, USA.

## Funding

This study was funded internally by Nanorx, Inc., NY, USA.

References

